# Deleting learning-induced dendritic spines disrupts the memory they encode

**DOI:** 10.64898/2026.08.18.745461

**Authors:** Hiranmay Joag, Patricio Opazo, Kenta M. Hagihara, Andreas Lüthi, Tobias Bonhoeffer

## Abstract

Long-term memory is widely thought to depend on activity-dependent synaptic plasticity and the structural remodeling that accompanies it, yet direct causal evidence that this remodeling is required for memory storage has been lacking. A case in point is the formation of dendritic spines: new spines appear following learning, but whether they constitute a physical substrate of memory remains unresolved. To address this, we developed the New Spine Elimination Tool (NSET), a chemical-genetic strategy, that ablates dendritic spines formed within a defined window of synaptic plasticity. NSET combines inducible expression of a degradable form of the actin-binding protein Drebrin with ligand-triggered proteasomal degradation. Because mainly nascent spines incorporate the degradable proteins into their cytoskeleton, ligand application eliminates them while sparing pre-existing ones. In hippocampal slice cultures, NSET eliminated recently formed spines without altering overall spine density or affecting pre-existing spines. Applied *in vivo* in the mouse basolateral amygdala, selective removal of learning-induced spines disrupted auditory fear memory, whereas consolidated memories and the capacity for new learning remained intact. These findings provide direct causal evidence that newly formed dendritic spines are required for long-term memory storage.

**One sentence summary:** Selectively eliminating new dendritic spines formed during learning disrupts the associated memory, demonstrating that newly formed spines are an essential component of long-term memory storage.

---

Long-term memory is widely believed to arise from activity-dependent modifications of synaptic function following learning, a phenomenon known as synaptic plasticity (*1, 2*). Synaptic plasticity underlying long-term memory is accompanied by pronounced and persistent structural remodeling of dendritic spines – tiny actin-rich protrusions that decorate neuronal dendrites and house the postsynaptic machinery of most excitatory synapses (*3–5*). This remodeling has been repeatedly observed as structural potentiation of pre-existing spines as well as the growth of entirely new dendritic spines both *in vitro* (*6–8*) and *in vivo* (*9–14*), reinforcing the hypothesis that structural synaptic plasticity represents an anatomical substrate of long-term memory storage (*4, 5*). Newly formed dendritic spines develop functional synapses (*15–17*) and show task-relevant functional responses (*18, 19*), however, whether these newly formed dendritic spines are necessary for long-term memory storage is unknown.

Here, we directly test this hypothesis using a newly developed chemical-genetic tool that selectively eliminates newly formed spines following the induction of synaptic plasticity or learning, while leaving pre-existing spines intact. This new spine elimination tool (NSET) employs a two-step strategy involving temporally controlled expression of a tagged postsynaptic actin-binding protein, Drebrin (*20, 21*), followed by its acute depletion using the dTAG system (*22*). We validated NSET in organotypic hippocampal slice cultures and subsequently demonstrated that its application *in vivo* in the mouse basolateral amygdala (BLA) selectively eliminates recently acquired fear memories while leaving previously acquired, consolidated memories intact. Together, these findings provide the first direct causal evidence that newly formed dendritic spines are required for long-term memory storage.

## Results

### A Drebrin-based molecular tool to eliminate dendritic spines

To test whether dendritic spines are causal substrates of memory storage, we developed a molecular tool that selectively eliminates newly formed dendritic spines while sparing pre-existing ones. Our reasoning was that selective ablation of only the spines generated during learning should result in loss of the corresponding memory and that such a result would provide direct evidence that dendritic spines serve as physical sites of information storage.

NSET was designed to target a cytoskeletal protein that is critical for spine stability, such that its acute removal or modification would trigger spine collapse. Moreover, temporal control over the tool had to be sufficiently precise to selectively label only those spines formed during the learning event. This in turn required rapid protein synthesis ensuring that newly formed spines would incorporate sufficient amounts of the modified protein to later serve as a reliable tag for degradation.

We first screened actin-binding proteins for their ability to destabilize dendritic spines upon acute inactivation (Fig. 1A). Candidate proteins fused to the phototoxic fluorescent protein KillerRed (*23*)were expressed in organotypic hippocampal slice cultures, and spine elimination was quantified following controlled illumination. Among all candidates, photo-destruction of Drebrin, an actin-stabilizing protein enriched in postsynaptic compartments(*21, 24*), produced the most robust and reproducible spine loss (Fig. 1A).

**Fig. 1.**
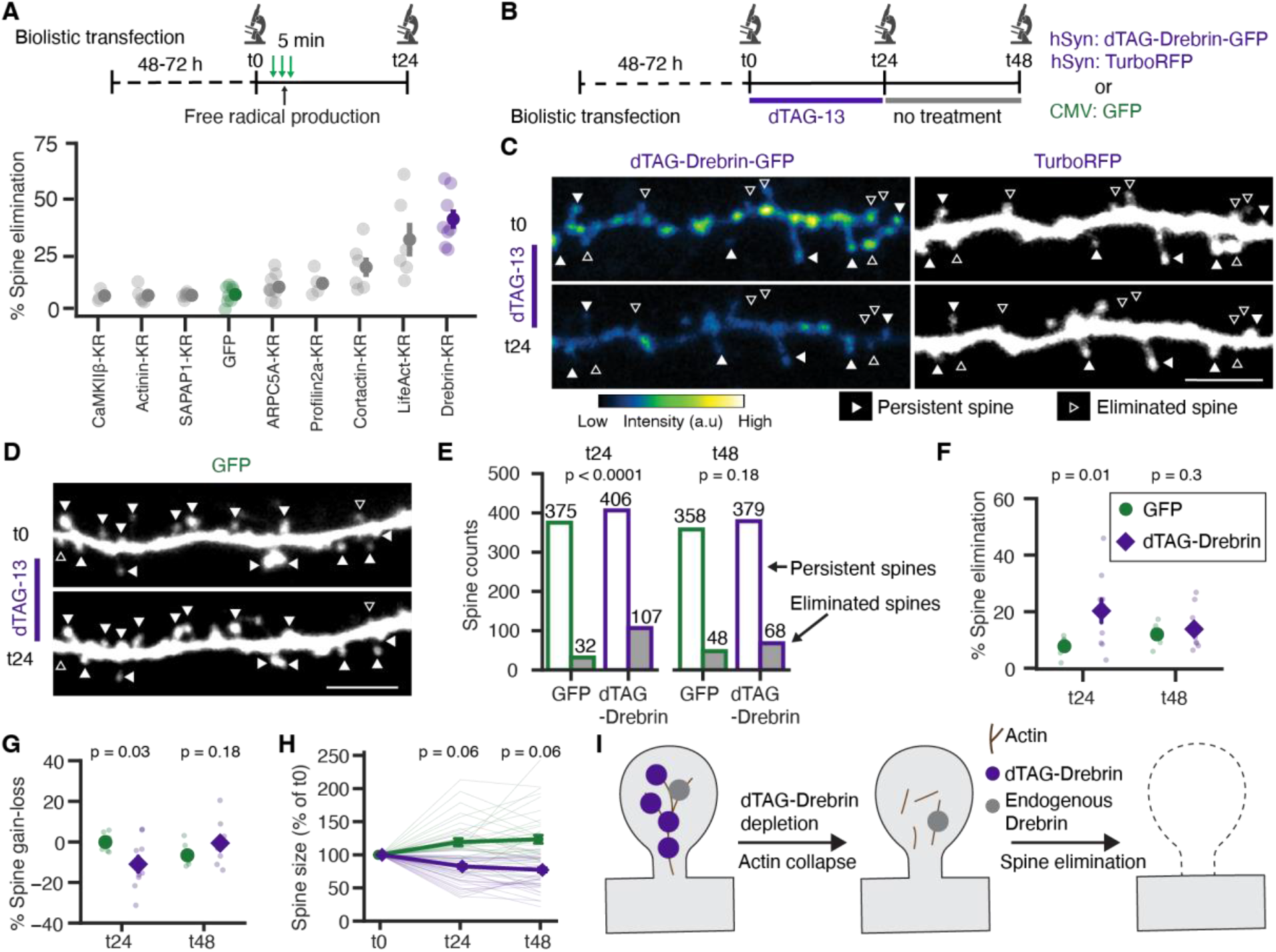
Acute dTAG-Drebrin depletion leads to spine elimination. **(A)** Experimental timeline (top) and percent dendritic spines eliminated 24 h after light-induced free radical production in neurons expressing the indicated actin-binding proteins tagged with KillerRed (KR). Green arrows indicate epifluorescence illumination used to induce free radical production. n = 3-9 neurons per actin-binding protein. (**B)** Experimental timeline (left) and constructs (right) used for the dTAG-Drebrin spine elimination assay. (**C, D)** Example images of dendritic segments expressing dTAG-Drebrin-GFP and TurboRFP (C) or GFP alone (D) before (t0) and after (t24) dTAG-13 treatment. Filled and open arrowheads indicate persistent and eliminated spines, respectively. Color bar represents arbitrary fluorescence units. Scale bars, 5 μm. (**E)** Chi-square analysis of persistent and eliminated dendritic spines from GFP- and dTAG-Drebrin-expressing neurons following dTAG-13 treatment (t24) and washout (t48). Raw spine counts are shown above the bars. (**F)** Branch-wise percentage spine elimination in 7 GFP- and 10 dTAG-Drebrin-expressing dendritic branches following dTAG-13 treatment (t24) and after washout (t48). (**G)** Branch-wise net spine change in the same branches as in (F), calculated as percent spine gain minus percentage spine elimination. (**H)** Spine size, normalized to baseline, before dTAG-13 treatment (t0), after treatment (t24), and following washout (t48). The numbers of spines tracked at t0, t24 and t48 are 133, 116 and 114 for GFP and 129, 74 and 60 for dTAG-Drebrin. (**I)** Proposed mechanism of spine elimination following acute dTAG-Drebrin depletion. All experiments were performed in rat organotypic hippocampal slice cultures. Quantitative data are presented as mean ± SEM except in (E), where total spine counts are shown. Chi-square analysis was used to compare spine fate proportions in (E). Statistical significance was assessed using linear mixed-effects models (LMM) (F, G) or bias-reduced generalized estimating equations (GEE) (H), followed by model-based contrasts with Holm-Bonferroni correction for multiple comparisons. No inferential statistics were performed for the screen in (A).

To enable reversible and ubiquitous control of Drebrin levels, we replaced KillerRed with the dTAG degradation system (*22*). Drebrin was fused to an inducible dTAG degron, which can be selectively targeted for proteasomal degradation by application of the small-molecule ligand dTAG-13. The ligand recruits the ubiquitously expressed E3 ligase Cereblon, thereby inducing ubiquitination and degradation of the tagged protein (*22*). We generated a degradable Drebrin construct (dTAG-Drebrin) and optimized its expression and depletion kinetics in hippocampal neurons (Fig. S1). We found that acute degradation of dTAG-Drebrin resulted in significant spine loss compared to controls following dTAG-13 treatment (Fig. 1B-I). Degradation of a truncated Drebrin isoform containing only the actin binding domain (dTAG-sDrebrin) (*25*), produced comparable effects (Fig. S2A-E), confirming that spine collapse resulted from destabilization of the actin cytoskeleton. Spine loss was accompanied by disappearance of PSD95Δ1.2, a PDZ domain deleted PSD-95(*26*), indicating synapse elimination (Fig. S2F-H).

### Selective elimination of newly formed spines *in vitro*

Because endogenous Drebrin has a long half-life (~3 days)(*27*), we reasoned that transiently expressed dTAG-Drebrin would predominantly accumulate in the cytoskeleton of nascent spines whereas pre-existing spines would mainly remain populated by endogenous Drebrin. Subsequent ligand-induced degradation of dTAG-Drebrin should therefore selectively eliminate newly formed spines (Fig. 2A).

**Fig. 2.**
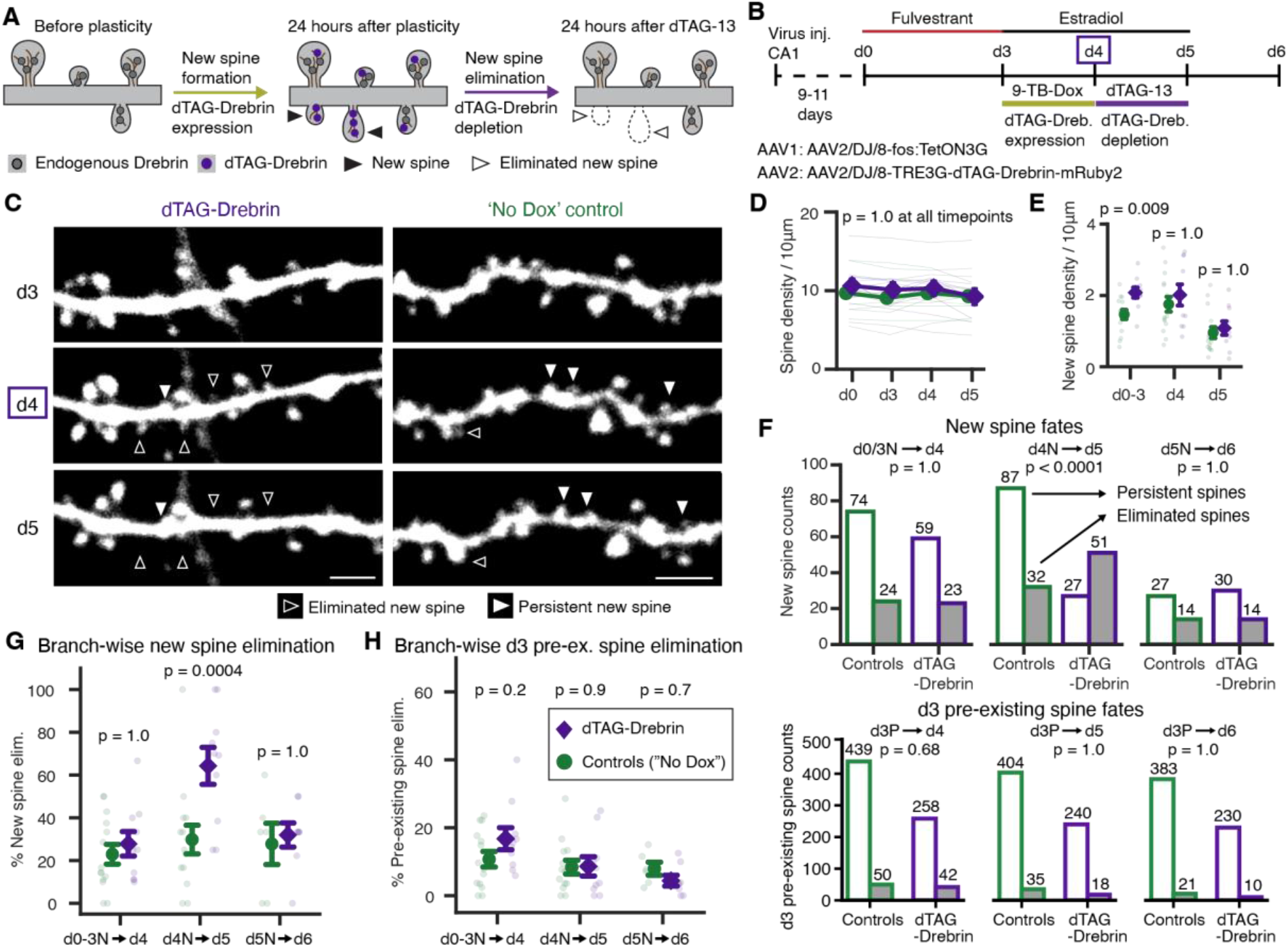
Selective elimination of newly formed dendritic spines *in vitro*. **(A)** Schematic illustrating the strategy for selective elimination of newly formed dendritic spines. dTAG-Drebrin is rapidly expressed following plasticity and is depleted 24 h later by dTAG-13 administration. **(B)** Experimental timeline (top) for selective new spine elimination in organotypic hippocampal slice cultures. dTAG-Drebrin expression was induced between d3 and d4 by 9-tert-Butyl Doxycycline (9-TB-Dox) administration. Following imaging on d4, dTAG-13 was administered to deplete dTAG-Drebrin, thereby selectively targeting d4 new spines. Viral constructs used for activity-dependent and doxycycline-dependent dTAG-Drebrin expression (bottom). (**C)** Example images of dendritic segments expressing the structural marker GFP in dTAG-Drebrin and “No Dox” control neurons before dTAG-Drebrin expression (d3), after activity-dependent dTAG-Drebrin expression (d4), and after dTAG-13 treatment (d5). Filled and open arrowheads indicate persistent and eliminated d4 new spines, respectively. Scale bar = 2.5 μm. (**D)** Branch-wise spine density normalized to dendritic path length (spines/10 μm) in 15 control and 10 dTAG-Drebrin branches throughout the imaging period. (**E)** Branch-wise density of new spines in the same branches shown in (D). The d0-3 time point represents spines that formed between d0 and d3. (**F)** Chi-square analysis of new (top) and d3 pre-existing spine fates (bottom). Spine populations were pooled across all branches and cells for each condition. Raw spine counts are shown above the bars. (**G)** Branch-wise elimination of newly formed spines in branches in which more than two new spines formed during the indicated interval. Controls: d0-3N→d4, n = 14 branches from 7 cells; d4N→d5, n = 14 branches from 6 cells; d5N→d6, n = 6 branches from 4 cells. dTAG-Drebrin group: d0-3N→d4, n = 10 branches from 6 cells; d4N→d5, n = 10 branches from 6 cells; d5N→d6, n = 8 branches from 6 cells. **(H)** Branch-wise elimination of d3 pre-existing spines in the same branches analyzed in (G). All experiments were performed in Thy1-GFP mouse organotypic hippocampal slice cultures. Graphical data are presented as mean ± SEM except in (F), where total spine counts are shown. Statistical significance was assessed using LMM (D, G) or GEE (E, H) followed by model-based contrasts with Holm-Bonferroni correction for multiple comparisons. Chi-square analysis was used to compare spine fate proportions in (F).

We first confirmed that we could indeed eliminate newly formed spines by driving dTAG-Drebrin expression with the activity dependent *fos* promoter(*28*) following Forskolin/Rolipram induced chemical long term potentiation (cLTP)(*29*) and degrading dTAG-Drebrin 24 hours following expression (Fig. S3). To selectively eliminate newly formed spines, we further restricted dTAG-Drebrin expression by placing it under control of the *fos* promoter as well as the doxycycline-inducible Tet System (*30*). To test this system we induced robust new spine formation by sequential Estrogen receptor degradation using Fulvestrant followed by Estradiol treatment in Thy-1 GFP mouse organotypic hippocampal slice cultures (*31, 32*). dTAG-Drebrin expression was restricted to a 24-hour window during Estradiol-induced spine formation (Fig. 2B). Subsequent application of dTAG-13 then indeed resulted in selective elimination of spines formed during the expression window, while pre-existing spines and total spine density remained unaltered (Fig. 2C-H, S4). We confirmed that similar selectivity was observed in an alternative Forskolin/Rolipram-induced cLTP paradigm (Fig. S5).

### Learning-induced spines are required for fear memory storage *in vivo*

We then used this tool to address the central, yet long-unresolved question: are learning-induced new spines necessary for long-term memory storage? We used auditory fear conditioning, a rapid form of associative learning dependent on the basolateral amygdala (BLA), which has been widely studied and where the locus of the synaptic changes is likely reasonably well confined (*33*). Using bilateral viral injections, we delivered either the full NSET construct (activity- and doxycycline-dependent dTAG-Drebrin-mRuby2) or control vectors lacking Drebrin to the BLA (Fig. S6 A-B). Guide cannulas were implanted above the BLA to allow subsequent local infusion of dTAG-13 (Fig. 3A). After recovery from the surgery and sufficient time for the expression of the virus, we performed three key experiments to address whether learning-induced new spines are necessary for memory storage and to ensure that the behavioral results were due to learning induced new spine removal and not due to off-target effects (Fig. 3B).

**Fig. 3.**
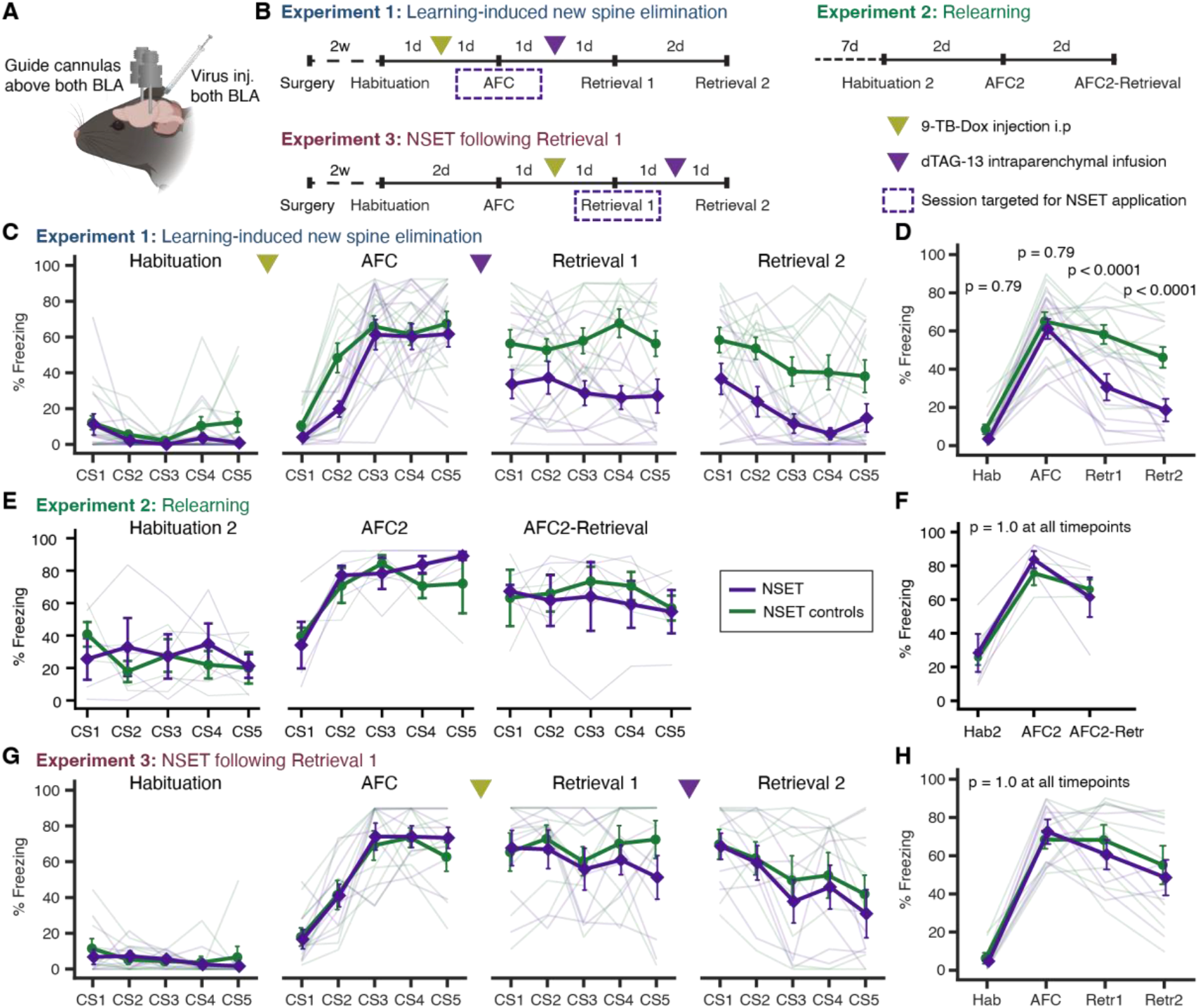
Learning-induced new spine elimination leads to memory loss. **(A)** Schematic illustrating bilateral viral injection and guide cannula placement above the basolateral amygdala (BLA). **(B)** Experimental timelines for the three *in vivo* paradigms shown in this figure. **(C, D)** Percent freezing during individual conditioned stimulus (CS) presentations (C) and mean freezing across CS presentations (D) during Habituation (Hab), Auditory Fear Conditioning (AFC), Retrieval 1 (Retr1), and Retrieval 2 (Retr2) in the learning-induced new spine elimination paradigm. Data are shown for 13 control and 12 NSET (New Spine Elimination Tool) mice. **(E, F)** Percent freezing during individual CS presentations (E) and mean freezing across CS presentations (F) during Habituation 2 (Hab2), Auditory Fear Conditioning 2 (AFC2), and Retrieval following AFC2 (AFC2-Retrieval) in the relearning paradigm. Data are shown for 3 control and 4 NSET mice. **(G, H)** Percent freezing during individual CS presentations (G) and mean freezing across CS presentations (H) during Habituation, Auditory Fear Conditioning, Retrieval 1, and Retrieval 2 in the NSET following Retrieval 1 paradigm. Data are shown for 8 control and 7 NSET mice. Graphical data are presented as individual mice together with group mean ± SEM. No inferential statistics were performed for panels (C), (E), and (G), which are included to visualize freezing behavior across individual conditioned stimulus (CS) presentations. Statistical significance was assessed using LMM followed by model-based contrasts with Holm-Bonferroni correction for multiple comparisons (D, F, H).

In the first experiment, mice underwent auditory fear conditioning (Fig. 3C-D), during which an initially neutral tone (conditioned stimulus, CS) was paired with a mild footshock (unconditioned stimulus, US). Both NSET and control groups showed comparable acquisition curves, with progressively increasing freezing behavior during CS exposure with each tone– shock pairing (Fig. 3C-D).

The following day, dTAG-13 was infused to degrade dTAG-Drebrin and eliminate learning-induced spines (Fig. 3C–D). When memory retrieval was tested 24 hours later, NSET-treated mice exhibited a marked reduction in freezing to the CS (Fig. 3C-D), suggesting that, indeed, deleting of the spines that were generated during the learning phase resulted in a loss of memory of the learned conjunction between tone and shock. Control animals, including those expressing inactive constructs or receiving vehicle instead of the dTAG-13, displayed robust memory retrieval (Fig. 3C-D, S6). Baseline behavior and locomotion were unaffected (Fig. S7D-E). Histological analysis confirmed correct bilateral viral expression and cannula placement in all behaviorally affected animals (Fig. S7F, S8). Animals with mistargeted injections or cannulas retained normal freezing responses (Fig. S7A-C).

In the second experiment, we controlled for the possibility that NSET may cause a generalized learning deficit. The same animals that had undergone experiment 1 were retrained on a second tone–shock association using a different auditory cue (Fig. 3B, E-F). NSET-treated mice acquired and retrieved this new association normally (Fig. Fig 3E-F, S7G-H), demonstrating that NSET did not generally affect learning.

Finally, in the third experiment, we tested the temporal specificity of NSET, by applying it after memory retrieval rather than during learning (Fig. 3B, G-H, Fig S9, 10). Even though the learning- and retrieval-activated cell populations are partially overlapping (*34–37*), degradation of dTAG-Drebrin had no effect on subsequent memory performance, in this experiment (Fig. 3G-H, Fig S9). This temporal dissociation demonstrates that memory loss results specifically from deletion of spines formed during learning, rather than from elimination or functional perturbation of spines at other times.

## Discussion

Together, these experiments demonstrate a causal and temporally specific requirement for learning-induced dendritic spines in memory storage. Using NSET, selective disruption of synaptic structures formed during a defined learning episode abolished the corresponding associative memory without affecting general learning capacity. These findings identify newly formed dendritic spines as essential physical substrates of memory.

While previous work has suggested a role of dendritic spine enlargement in memory storage (*9*), our study addresses a fundamentally different question that has remained unresolved: whether the formation of new spines is necessary for long-term memory storage. Thus, rather than examining how structural changes in existing spines contribute to memory, our experiments directly test the causal requirement for learning-induced spine formation using a specific and easily applicable tool. We hope, this will facilitate its adoption across a broader range of experimental systems and enable answering questions about spine plasticity that have thus far been difficult to address with existing approaches.

Another important consideration is that NSET, as it targets the cytoskeleton, may also affect synaptic function, however again only affecting synapses that have been generated at the time of memory-acquisition. Acute degradation of Drebrin is expected to destabilize the postsynaptic actin cytoskeleton, potentially disrupting postsynaptic density organization and glutamate receptor localization (*21, 24, 38*) before overt spine collapse. Thus, memory loss may result from functional disruption of newly formed excitatory synapses in addition to, or preceding, their structural elimination.

The substantial, but incomplete, memory impairment (Fig 3C-D) can be explained by a limited efficacy of NSET (be it with respect to reaching the entire BLA or affecting all spines), but it is likely also due to the fact that other processes such as structural potentiation of pre-existing spines (*9, 10, 12, 39, 40*), changes in intrinsic excitability (*41–44*), axonal plasticity (*45*) plasticity of inhibitory circuits (*46*), and BLA-independent processes (*33*) play important roles in memory persistence.

Our findings extend beyond fear conditioning and the BLA. Structural plasticity is a widespread feature of cortical and hippocampal circuits, and NSET provides a generalizable approach to test the causal contribution of experience-dependent spine formation across brain regions and behavioral paradigms. Because NSET relies on genetically encoded and pharmacologically inducible components, it offers precise anatomical, cellular and temporal control providing a versatile platform for dissecting the contribution of structural plasticity to learning, memory and circuit remodeling.

Conceptually, our results bridge decades of correlational evidence linking new spine formation to long-term memory. They support the idea that transient synaptic signals are converted into stable memories through enduring structural modifications in neuronal connectivity. By demonstrating that selective removal of learning-induced dendritic spines abolishes a specific memory, this work provides direct experimental evidence that memory is embodied in the connectivity and structure of neuronal circuits.

## Supporting information

Joag et al Supplementary Materials

## Acknowledgements

We thank Volker Staiger, Claudia Huber, Dominik Lindner, Frank Voss, Max Sperling, Tobias Eichlisberger, Nigel Whittle and Sigrid Müller for technical support, Sascha-Alexander Heye for spine annotation, Facility for Advanced Imaging at the FMI for Biomedical Research for imaging assistance, Joel Bauer and Nicolas Karalis for help with coding, Behnam Nabet, Craig Crews and Brenda Schulman for advice on targeted protein degradation, and all members of the Lüthi and Bonhoeffer laboratories for discussions.

## Author contributions

H.J., A.L. and T.B. designed the project. H.J. performed all experiments and imaging data analysis. H.J. and K.M.H. tested the viral constructs *in vivo*. P.O. performed the KillerRed screen. All authors contributed to the experimental design, interpretation of the data and wrote the manuscript.

## Data and materials availability

All data and analyses necessary to understand and assess the conclusions of the manuscript are presented in the main text and in the supplementary materials. Processed data and code relating to this paper will be deposited in a publicly available repository upon publication. Plasmids used in this manuscript will be made available through Addgene.

## Supplementary Materials

Materials and Methods fig. S1 to S10

