## Supplementary material for "Deleting learning-induced dendritic spines disrupts the memory they encode": Joag et al Supplementary Materials

**This PDF file includes:**

Materials and Methods
Table S1, Figs. S1 to S10

### Materials and Methods

#### Experimental design and blinding

No statistical methods were used to pre-determine sample sizes for any of the experiments. Organotypic cultures were made from male and female pups and randomly allocated to experimental groups. Dendritic spine annotation was performed by investigators blinded to the experimental groups. For *in vivo* experiments, male and female mice were randomly allocated to the experimental groups. The experimenter was not blinded during behavioral testing or freezing analysis; however, freezing was quantified automatically using a fixed threshold without manual scoring or post hoc adjustment. Histological classification was performed by investigators blinded to experimental group and behavioral outcomes.

#### *In vitro* experiments

Plasmids

Plasmids used and generated in this study are listed in table S1. KillerRed fusion constructs and hSyn-PSD95-Δ1.2-GFP were generated through restriction enzyme cloning. All other plasmids were generated using Gibson assembly (*1*). All inserts were sequence-verified. Complete maps and sequences of newly generated plasmids will be deposited on Addgene.

Organotypic hippocampal slice cultures (OHSCs)

OHSCs were prepared from 5-9 day old pups in compliance with Max Planck Society guidelines and Regierung von Oberbayern regulations. 400 µm thick OHSCs were prepared from Wistar rats (Figs. 1 and S1-3), Thy1-GFP (*2*) (Figs. 2 and S4) or C57BL/6 (Fig S5) mice as previously described (*3*) . Culture medium was supplemented with penicilin-streptomycin (Omnilaboratories A8943,0100). Imaging experiments were performed on cultures maintained *in vitro* for 2 - 4 weeks.

Biolistic transfection

Cultures were biolistically transfected 2–3 days before the first experimental time point using a Helios Gene Gun (Bio-Rad, 1652432), 1.6-µm gold particles (Bio-Rad, 1652264), helium at 180 psi, and 6–8 µg DNA per construct. Biolistic transfection was used for Figs. 1, S1A–E, S2 and S3.

Virus injections *in vitro*

AAV transduction was used for experiments in Fig S1F, Fig 2, S4 and S5. Viruses were injected into CA1 1–4 days after culture preparation using borosilicate capillaries (Harvard Apparatus, GC150F-10) connected to a Pneumatic PicoPump (World Precision Instruments, PV820). Brief 20-psi pulses generated droplets approximately 80 µm in diameter, and one or two droplets were injected per culture. Imaging began 9–11 days later. AAV2/DJ8-fos-TetOn3G-WPRE ($3.56 \times 10^{12}$ GC/ml; Vectorbuilder) and AAV2/DJ8-TRE3G-dTAG-Drebrin-mRuby2-WPRE ($1.24 \times 10^{13}$ GC/ml; Vectorbuilder) were injected into rat, Thy1-GFP or C57BL/6 mouse OHSCs. For sparse labeling in C57BL/6 cultures (Fig. S5), AAV2/9-CaMKII(0.4)-Cre-SV40 ($9.62 \times 10^{9}$ GC/ml; University of Pennsylvania Vector Core) and AAV2/1-CAG-FLEX-EGFP-WPRE ($3.19 \times 10^{12}$ GC/ml; University of Pennsylvania Vector Core) were additionally coinjected. Viruses were diluted in cortex buffer containing 125 mM NaCl, 5 mM KCl, 10 mM D-glucose, 10 mM HEPES, 2 mM $CaCl_2$, and 2 mM $MgSO_4$ (pH 7.4).

Two-Photon imaging

Chronic spine imaging was performed with a custom-built two-photon laser-scanning microscope described previously (*4*) (Olympus 60×/0.9 NA objective, Mai Tai Ti:Sapphire laser - Mai Tai Spectra Physics HP, Hamamatsu PMTs). 1024 * 1024-pixel images were obtained with a 2-µs pixel dwell time and 1-µm z-steps. Excitation wavelengths were 920 nm for imaging GFP and 1020 nm for mRuby2 or TurboRFP. Laser intensity at the objective was 5-10 mW. The same dendritic branches were tracked across imaging sessions using baseline reference images and local anatomical landmarks. OHSCs were continuously perfused during the imaging session with carbogenated (95% v/v $O_2$ and 5% v/v $CO_2$) artificial cerebrospinal fluid (ACSF) consisting of 125 mM NaCl, 2.5 mM KCl, 1.25 mM $NaH_2PO_4$, 1 mM $MgCl_2$ hexahydrate, 26 mM $NaHCO_3$, 2 mM $CaCl_2$ dihydrate, 25 mM D-glucose, and 200 µM Trolox in distilled water. From 1 day before the first imaging session until the end of the experiment, 200 µM Trolox (Sigma-Aldrich Cat. No.238813) was added to the culture medium. Trolox was excluded from the culture medium and ACSF for the KillerRed screen in Fig 1A.

Epifluorescence screens

For Figs. S1A–E, OHSCs were biolistically cotransfected with fos-TetOn3G and TRE3G-dependent constructs encoding N- or C-terminal dTAG fusions. For Fig. S1F, cultures were transduced with the AAVs described above. Expression was induced with 9-TB-Dox (1 µg/ml, Echelon Biosciences Cat. No. B-0801); added 2 h before bicuculline (30 µM; Tocris Cat. No. 2503), which was applied for 45 min. Cultures were washed, maintained in 9-TB-Dox for 24 h, and transferred to doxycycline-free medium for 24 h before baseline imaging.

After baseline imaging, HaloPROTAC3 (a kind gift from Craig Crews(*5*)), dTAG-13 (kind gift from Behnam Nabet and also purchased from Tocris Cat. No. 6605) (*6*) or dTAG-v1 (kind gift from Behnam Nabet) (*7*), was added at the concentrations indicated in Fig. S1, with matched DMSO controls. The same regions were imaged after 24 h. For Fig. S1F, compounds were washed out and cultures were imaged again 24 h later.

The same regions were imaged at each time point using identical settings. Cell ROIs were drawn in ImageJ and mean fluorescence was measured after background subtraction. Relative change was calculated as percent change in fluorescence relative to baseline.

KillerRed screen

After baseline 2-photon imaging, a selected dendritic region was illuminated continuously for 5 min through the Olympus 60×/0.9 NA objective using the green epifluorescence filter set and an Olympus U-LH100HG 100-W mercury lamp operated at maximum output. The same dendritic branches were imaged again 24 h later.

Nonselective spine elimination

After baseline imaging of one or two branches per cell, cultures were treated with dTAG-13 (0.5 µM) for 24 h and reimaged. dTAG-13 was then washed out, and a final imaging session was performed 24 h later (Figs. 1 and S2).

New spine elimination experiments (Fig 2, S3-5)

For forskolin/rolipram-induced structural plasticity experiments (Figs. S3 and S5), cultures were treated with forskolin (25 µM) and rolipram (2.5 µM) in $Mg^{2+}$-free ACSF for 17 min after baseline imaging. In Fig. S5, 9-TB-Dox (1 µg/ml) was included in the ACSF during induction and subsequently in the culture medium for 24 h.

For estradiol-induced structural plasticity experiments (Figs. 2 and S4), fulvestrant (1 µM) was added to the culture medium from d0 to d3. After imaging on d3, fulvestrant was washed out and β-estradiol (1 µM) was added from d3 to d5. 9-TB-Dox (1 µg/ml) was applied together with β-estradiol from d3 to d4.

Spine analysis

A total of 17,320 spines from 119 dendritic branches, belonging to 61 cells, were tracked across 2-5 imaging sessions. Spine counting and matching were performed in ImageJ using a custom-modified ROI manager tool and as per well-established criteria(*8*, *9*). Each plotted datapoint represents one dendritic branch.

Spine density was defined as the spines per 10 µm dendrite. Percent spine loss was calculated as the number of spines lost, relative to the number of spines present at the previous timepoint. Spine turnover was calculated as the percentage of spines gained minus the percentage of spines lost for that branch. For percent new and pre-existing spine elimination, only those branches were included in the analysis, where >2 new spines grew following structural plasticity.

For PSD-95Δ1.2 and dTAG-Drebrin colocalization analysis, protrusions were classified as spines if they contained clear enrichment of either PSD-95Δ1.2-GFP or dTAG-Drebrin-mRuby2. Similarly, a shaft synapse was defined as a clear enrichment of either marker on the dendritic shaft. The percentage of PSD-95Δ1.2-positive structures containing dTAG-Drebrin and the percentage of dTAG-Drebrin-positive structures containing PSD-95Δ1.2 were calculated.

Spine size was estimated as the ratio of the structural marker's fluorescence intensity in a ROI drawn around a spine to an ROI drawn in the dendritic segment at the base of the spine(*10*) . All measurements were performed on noise-subtracted, averaged z-projections. For all longitudinal analyses, these size ratios were normalized to their baseline (t0) value. Within this longitudinal dataset, spines were further classified for subpopulation analyses: 'enlarged spines' were defined as those that underwent a ≥25% increase in size compared to the previous day, while 'non-enlarged spines' were all other persistent spines. For comparisons of overall size distributions (Fig S4B), raw, non-normalized ratios were used.

### ***In vivo* experiments**

#### Mice

All animal protocols were in accordance with institutional guidelines of the Friedrich Miescher Institute (FMI) for Biomedical Research and were approved by the Cantonal Veterinary authorities of Basel-Stadt. C57BL/6J (C57BL/6JRccHsd; Envigo) mice were used in the study. Mice were singly housed in cages enriched with a running wheel, a cardboard or glass tunnel and straw for at least 2 weeks before auditory fear conditioning. Mice had *ad libitum* access to food and water and were maintained on a 12-hour light/dark cycle. Experiments were performed during the light phase.

#### Stereotaxic surgeries

8–10-week-old mice were anesthetized with isoflurane (3-5% for induction, 1-2% maintenance, Attane, Provet), fixed in a stereotaxic apparatus (Model 1900, Kopf Instruments) and 300 nl viral mixture per hemisphere was loaded into a glass pipette and injected into the BLA (Viral mixture consisted of AAV2/DJ8 fos-flex-d2TetOn3G: 3.6 × $10^{12}$ GC/ml; Vectorbuilder, AAV2/9 CaMKII (0.4)-Cre: 1.5 × $10^{12}$ GC/ml; University of Pennsylvania Vector Core in addition to AAV2/DJ8 TRE3G-dTAG-Drebrin-mRuby2: 2.5 × $10^{13}$ GC/ml; Vectorbuilder or AAV2/DJ8 TRE3G-dTAG-mRuby2: 5.4 × $10^{13}$ GC/ml; Vectorbuilder) with a picospritzer (Picospritzer III, Parker Hannifin Corporation) at the following coordinates: AP (from bregma): -1.55, ML (from bregma): ± 3.4, DV (from pia): -4.4 (200 nl) and -4.2 (100 nl). Following virus injection, guide cannulas (RWD 26G, Cat 6212) were inserted up to DV -3.4. The guide cannulas were affixed to the skull using UV light curable glue (Henkel, Loctite 4305), following which the skull was lightly scratched, covered with Optibond™ followed by dental cement (Paladur; Kulzer). Cap cannulas (RWD 26G, Cat 6212) were then inserted and screwed into the guide cannulas. Pre- and postoperative care and analgesia was provided as described in (*11*).

#### Drugs *in vivo*

For *in vivo* experiments, 25 µg/g 9-TB-Doxycycline was freshly dissolved in normal saline and injected intraperitoneally one day before Auditory Fear Conditioning (Fig 3, S6-8) or Retrieval 1 (Fig 3, S9-10). For dTAG-13 infusions, dTAG-13 was freshly dissolved in DMSO to create a 47.65 mM dTAG-13 solution. 1 µl of dTAG-13 solution was diluted in 19 ul of 20% Solutol in saline. 0.5 µl of this formulated solution (final molarity 2.385 mM) was infused through an injection cannula (RWD 26G, Cat 6212) that fit snugly into the guide cannula and protruded 1mm beyond the guide cannula, resulting in an infusion at DV -4.4. Vehicle-control mice received the same DMSO/Solutol/saline formulation without dTAG-13, using the same volume, infusion rate, and procedure. Infusion was performed at 0.3 µl/min with an infusion pump (New Era pump systems NE-300, WPI Flex PE tubing 504278) in mice briefly anesthetized using isoflurane. The injection cannula was kept in place for an additional 2 minutes following infusion and then withdrawn. No behavioral experiments were performed on the day of infusion or intraperitoneal (i.p.) injection.

Behavioral paradigms

Mice were handled and habituated to the experimenter for 5 days before commencing behavioral experiments. Behavioral experiments were conducted in two custom-built chambers equipped with overhead speakers for auditory stimulus delivery and cameras (Stingray, Allied Vision) for behavioral tracking. Auditory stimuli were generated using a System 3 RP2.1 real-time processor and SA1 stereo amplifier controlled by RPvdsEx software (Tucker-Davis Technologies). Behavioral protocols were controlled using TTL pulses generated with Radiant software (Plexon), and all TTL signals were recorded with PlexControl software (Plexon) to synchronize stimulus presentation, behavioral events, and video acquisition.

Habituation and retrieval were performed in a clear round Plexiglas arena with a smooth floor in the presence of 1% acetic acid (Context B). Mice were presented with five conditioned stimuli (CS; 7.5 kHz, 250-ms pips at 1 Hz, 20 s total duration, 75 dB sound pressure level, 60–90 s inter-trial intervals). Auditory fear conditioning (AFC) was performed in a square Plexiglas chamber with a grid floor for footshock delivery (Coulbourn Instruments) in the presence of 70% ethanol (Context A). During AFC, each CS co-terminated with a 1-s, 0.6-mA AC footshock.

For the relearning experiment, habituation and retrieval were performed in Context B', consisting of a large clear square Plexiglas arena scented with 1% isoamyl acetate. Mice were presented with five CS (white noise, 250-ms pips at 1 Hz, 20 s total duration, 75 dB sound pressure level, 60–90 s inter-trial intervals). AFC was performed in Context A', consisting of a round Plexiglas chamber with white walls, scented with 1% citral and equipped with a shock grid distinct from that used during the initial fear-conditioning session.

Freezing was quantified automatically using custom-written code and defined as movement below a fixed threshold (0.15) for at least 2 s. The same threshold was applied to all mice and sessions without manual adjustment or post hoc classification. Percent time spent freezing during each 20-s CS presentation or corresponding averaged 20-s bins throughout the 2 minute baseline period is reported.

Histology

At the end of behavioral experiments, mice were deeply anesthetized by administering 250 mg/kg ketamine and 2.5 mg/kg medetomidine solution intraperitoneally and transcardially perfused with 0.1M PBS for 2 minutes followed by 4% paraformaldehyde (PFA) in PBS for 10 minutes. Brains were post-fixed in 4% PFA for 24-48 h at 4 °C, and cut into 100 µm sections using a Leica VT1200S vibratome, mounted with Fluoromount-G containing DAPI, and imaged using a Zeiss Axio Scan.Z1 slide scanner with a Zeiss Fluar 5×/0.25 objective.

Histological analyses were performed blinded to the behavioral data. Mice were classified as NSET when viral expression was observed bilaterally throughout the BLA and both cannula infusion sites were located within the BLA. Mice with incomplete viral coverage of one or both BLAs or with misplaced cannula infusion sites were classified as NSET-mistargeted. Two animals were excluded from the analysis post hoc: one because tissue damage during histological processing prevented reliable histological assessment and one because one cannula track terminated at the intersection of the posterior BLA and the lateral ventricle, precluding reliable determination of whether the infusion reached the target region.

The histological reconstructions shown in Figs. S8 and S10 summarize viral expression and cannula infusion sites for all mice included in the behavioral experiments. Approximate viral spread and cannula infusion sites were manually mapped from serial coronal sections onto a single corresponding coronal plate at bregma −1.84 mm from the Paxinos and Franklin mouse brain atlas(*12*).

Activity- and doxycycline-dependent viral expression (Fig. S6) was verified post hoc by confocal imaging of coronal sections using a spinning-disk microscope equipped with a W1 scan head with 50-µm pinholes (Yokogawa) and a Plan-Apochromat 10×/0.45 air objective with a 1.5× Optovar (Zeiss). Images were acquired using VisiView 6.0 software (Visitron).

**Illustrations and example-image preparation**

Experimental timelines and illustrations were created using Adobe Illustrator and BioRender.com and exported under BioRender publication licenses, where applicable. Example images of dendritic segments were generated from average z-projections and median-filtered in ImageJ using a 2-pixel radius. For comparisons of fluorescence across time points – incuding dendritic branches in Fig 1, S2 and S3; cellular fluorescence in Fig S1; and activity- and doxycycline- dependent dTAG-Drebrin expression in the BLA *in vivo* in Fig S6 - corresponding z-projections or single plane images (Fig S1) were combined into a single stack and to ensure identical background and fluorescence scaling across images. Images of structural markers and the example histological images in Figs. S7 and S9 were adjusted individually, with brightness and contrast adjustments applied globally across each image. DAPI and dTAG-Drebrin-mRuby2 fluorescence in the histological images were displayed using blue and white lookup tables, respectively. Spine markers, cannula locations, and approximate BLA boundaries were added as overlays in Adobe Illustrator. Image processing was performed only for visualization and did not affect quantitative analyses.

**Statistical analysis**

Statistical analyses were performed in Python 3.13.9 using statsmodels 0.14.5 and SciPy 1.16.3. NumPy 2.3.5 and pandas 2.3.3 were used for data processing, and Matplotlib 3.10.6 and seaborn 0.13.2 for visualization. Unless otherwise stated, data are shown as mean ± SEM. All tests were two-sided, with $p < 0.05$ considered statistically significant.

Linear mixed-effects models (LMMs), fitted by restricted maximum likelihood, were the primary method used to analyze continuous outcomes containing repeated or nested observations. For *in vitro* experiments, experimental group, time point, and their interaction were included as fixed effects where applicable, with cell identity as a random intercept. Measurements from individual dendritic branches were treated as separate observations, while the cell-level random effect accounted for repeated measurements and multiple branches from the same cell. For behavioral experiments, mean CS-induced freezing was modeled as a function of experimental group, behavioral session, and their interaction, with mouse identity included as a random intercept.

LMM residuals were assessed using Shapiro–Wilk tests and quantile–quantile plots. When residuals deviated from normality, the data were instead analyzed using a Gaussian generalized estimating equation (GEE) model. These models used the same fixed effects as the corresponding LMM, with cell or mouse identity as the clustering variable, an exchangeable correlation structure, and a bias-reduced covariance estimator.

Planned model-based contrasts compared experimental groups at each time point or behavioral session. Contrasts were evaluated using two-sided Wald z tests, and p values were adjusted for multiple comparisons within each analysis using the Holm method. These Holm-adjusted P values are reported in the figures.

For spine-fate analyses, persistent and eliminated spines were pooled across the relevant branches and cells within each experimental group. The resulting proportions were compared using Pearson's chi-square tests on 2 × 2 contingency tables with Yates' continuity correction. P values from related spine-fate comparisons within each figure were adjusted using the Holm method. For the empirical distributions in Fig. S4B, spine sizes were compared between groups at each time point using two-sample Kolmogorov–Smirnov tests, with Holm correction across time points.

### Table S1 Plasmids used in the study

| Construct | Full construct name | Figure Reference | Backbone | Insert(s) |
|---|---|---|---|---|
| **1. CaMKIIβ-KR** | CMV- CaMKIIβ-KillerRed | Fig. 1A | pKillerRed-C (Evrogen FP961) | CaMKIIbeta from GFP-CaMKIIbeta (Addgene #21227) (*13*) |
| **2. Actinin-KR** | CMV- α-actinin-1-KillerRed | Fig. 1A | pKillerRed-N (Evrogen FP962) | α-actinin-1 from GFP-α-actinin-1 (Addgene #11908) (*14*) |
| **3. SAPAP1-KR** | CMV-SAPAP1-KillerRed | Fig. 1A | pKillerRed-C (Evrogen FP961) | SAPAP1 from Myc-SAPAP1 (Addgene #40215) (*15*) |
| **4. ArpC5A-KR** | CMV-ArpC5A-KillerRed | Fig. 1A | pKillerRed-C (Evrogen FP961) | ArpC5A from GFP-ArpC5A (gift from Gregory Giannone) (*16*) |
| **5. Profilin2a-KR** | CMV-Profilin-2a-KillerRed | Fig. 1A | pKillerRed-N (Evrogen FP962) | Profilin-2a from Profilin-2a-EGFP-N1 (gift from Walter Witke) (*17*) |
| **6. Cortactin-KR** | CMV-Cortactin-KillerRed | Fig. 1A | pKillerRed-C (Evrogen FP961) | Cortactin from Cortactin-pmCherry-C1 (Addgene #27676) (*18*) |
| **7. LifeAct-KR** | CMV-LifeAct-KillerRed | Fig. 1A | pKillerRed-N (Evrogen FP962) | LifeAct from LifeAct-GFP (gift from Roland Wedlich-Söldner) (*19*) |
| **8. Drebrin-KR** | CMV-Drebrin A-KillerRed | Fig. 1A | pKillerRed-C (Evrogen FP961) | Drebrin A from pEGFP-C1-Drebrin A (gift from Tomoaki Shirao) (*20*) |
| **9. GFP** | pEGFP-N1 (Clontech 6085-1) | Fig. 1B-H; Figs. S2 and S3 | Not applicable. | Not applicable. |
| **10. hSyn-TurboRFP** | pENN-AAV-hSyn-TurboRFP-WPRE-RBG (Addgene plasmid #105552) | Fig. 1B-C; Figs. S2A-E, S3 | Not applicable. | Not applicable. |
| **11. hSyn-dTAG-Drebrin-GFP** | pAAV-hSyn-dTAG-Drebrin A-GFP-WPRE | Fig. 1B-I; Fig. S2A-E | Derived from pAAV-EF1alpha-F-FLEX-Kir2.1-T2A-tdTomato (Addgene #60661) (*21*) | FKBP12(F36V) dTAG from pLEX_305-N-dTAG (Addgene #91797) (*6*); Drebrin A from pEGFP-C1-Drebrin A (*20*); GFP from pEGFP-N1 (Clontech 6085-1). |
| **12. hSyn-dTAG-sDrebrin-GFP** | pAAV-hSyn-dTAG-s-Drebrin A-GFP-WPRE | Fig. S2A-E | pAAV-hSyn-dTAG-Drebrin A-GFP-WPRE | Drebrin A excised and s-Drebrin A sequence from (*22*) amplified and inserted between dTAG and GFP. |
| **13. fos-dTAG-Drebrin-GFP** | fos-dTAG-Drebrin A-GFP | Fig. S3 | AAV2-ITR backbone (VectorBuilder VB200224) | dTAG-Drebrin-GFP excised from hSyn-dTAG-Drebrin-GFP plasmid above. |
| **14. Tet-On Advanced** | pTet-On Advanced (Clontech 630930) | Fig. S2F-H | Not applicable. | Not applicable. |

| | | | | |
|---|---|---|---|---|
| **15. PSD-95Δ1.2-GFP** | hSyn-PSD95-Δ1.2-GFP | Fig. S2F-H | Derived from pAAV-EF1alpha-F-FLEX-Kir2.1-T2A-tdTomato (Addgene #60661) (*21*) | PSD-95Δ1.2-GFP from pCAFNF-PSD-95Δ1.2-GFP (Addgene #125581) (*23*) |
| **16. fos-TetOn3G** | pAAV-fos-TetOn3G-WPRE | Fig. 2; Figs. S4 and S5 | Derived from pAAV-CaMKIIalpha-hM4D(Gi)-mCherry-WPRE (Addgene #50477; Bryan Roth) | c-fos promoter from pAAV-cFos-tTA-pA (Addgene #66794) (*24*) TetOn3G from pLVX-Tet3G-blasticidin (Addgene #128061; Oskar Laur) |
| **17. TRE3G-dTAG-Drebrin-mRuby2** | pAAV-TRE3G-dTAG-Drebrin A-mRuby2-WPRE | Figs. 2 and 3; Figs. S2F-H and S4-S10 | AAV2-ITR backbone (VectorBuilder VB200224) | TRE3G from pLVX-TRE3G-BFP (Addgene #128070; Oskar Laur); dTAG from pLEX_305-N-dTAG (Addgene #91797) (*6*) Drebrin A from pEGFP-C1-Drebrin A (*20*); mRuby2 from Addgene #68717 (*25*) |
| **18. TRE3G-dTAG-sDrebrin-mRuby2** | pAAV-TRE3G-dTAG-s-Drebrin A-mRuby2-WPRE | Fig. S1 | AAV2-ITR backbone (VectorBuilder VB200224) | TRE3G and dTAG as above; s-Drebrin A derived from hSyn-dTAG-sDrebrin-GFP, mRuby2 from Addgene #68717. (*25*) |
| **19. TRE3G-mRuby2-Drebrin-dTAG** | pAAV-TRE3G-mRuby2-Drebrin A-dTAG-WPRE | Fig. S1 | AAV2-ITR backbone (VectorBuilder VB200224) | Assembled from TRE3G-dTAG-Drebrin-mRuby2 plasmid described above. |
| **20. TRE3G-mRuby2-sDrebrin-dTAG** | pAAV-TRE3G-mRuby2-s-Drebrin A-dTAG-WPRE | Fig. S1 | AAV2-ITR backbone (VectorBuilder VB200224) | Assembled from the TRE3G-dTAG-sDrebrin-mRuby2 plasmid described above. |
| **21. TRE3G-HaloTag7-sDrebrin-mRuby2** | pAAV-TRE3G-HaloTag7-s-Drebrin A-mRuby2-WPRE | Fig. S1 | AAV2-ITR backbone (VectorBuilder VB200224) | s-Drebrin-mRuby2 from plasmid 18. Above; HaloTag7 from pET28a-HaloTag7-SnoopLigase (Addgene #105627) (*26*) |
| **22. fos-flex-d2TetOn3G** | pAAV-fos-flex-d2TetOn3G-WPRE | Fig. 3; Figs. S6-S10 | AAV2-ITR backbone (VectorBuilder VB200224) | TetOn3G derived from fos-TetOn3G plasmid described above; d2 degradation sequence/linker from Addgene #63931) (*27*) |
| **23. TRE3G-dTAG-mRuby2** | pAAV-TRE3G-dTAG-mRuby2-WPRE | Fig. 3; Figs. S6-S10 | AAV2-ITR backbone (VectorBuilder VB200224) | Drebrin excised from the above described plasmid TRE3G-dTAG-Drebrin-mRuby2 to create this insert. |

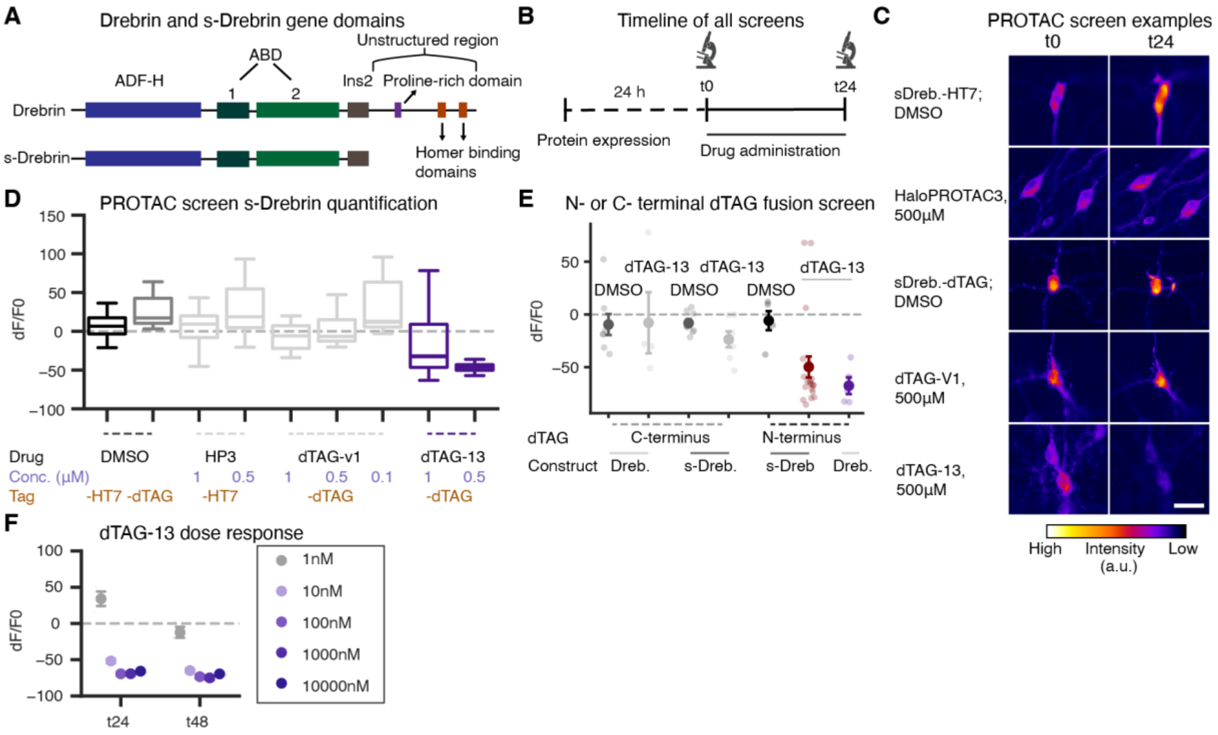


**Fig. S1. Development and characterization of the dTAG-Drebrin and s-Drebrin degradation strategy.**

**(A)** Schematic showing the major domains of Drebrin A and short Drebrin A (s-Drebrin A), referred to throughout the manuscript as Drebrin and s-Drebrin, respectively. **(B)** Experimental timeline for the fluorescence-based degradation screens shown in this figure. Panel F additionally includes a t48 imaging time point. **(C)** Example images from the PROTAC screen showing cellular fluorescence of tagged s-Drebrin (s-Dreb.) before (t0) and after 24 h of treatment (t24). Scale bar = 5.5 μm. **(D)** Change in tagged s-Drebrin cellular fluorescence relative to baseline after 24 h of treatment with the indicated PROTACs. Drug, concentration, and tag configuration are indicated below the x-axis. $n$ = 8-32 cells per drug condition. **(E)** Change in cellular fluorescence after 24 h of dTAG-13 treatment in cells expressing N- or C-terminally tagged Drebrin (Dreb.) or s-Drebrin (s-Dreb.) constructs. $n$ = 8-22 cells per condition. **(F)** Change in bulk fluorescence following treatment with increasing concentrations of dTAG-13 in cultures expressing activity- and doxycycline-dependent dTAG-Drebrin. dTAG-13 was applied from t0 to t24, followed by washout from t24 to t48. $n$ = 3-4 cultures per dose.

All experiments were performed in rat organotypic hippocampal slice cultures. In (D), boxplots show the median and interquartile range, and whiskers extend to 1.5 times the interquartile range. Data in (E) and (F) are shown as mean ± SEM. No inferential statistics were performed.

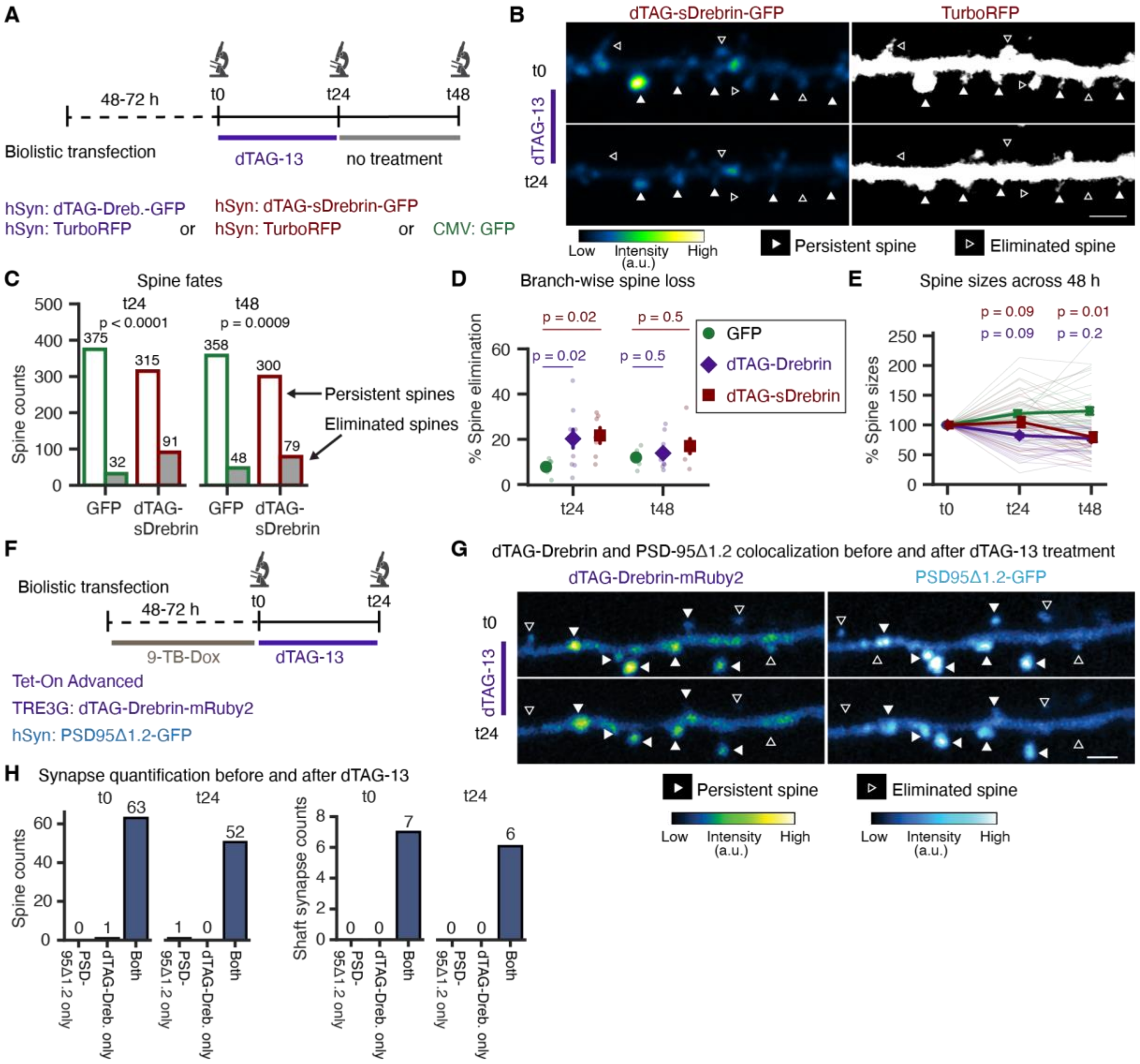


**Fig. S2. Acute dTAG-Drebrin or dTAG-sDrebrin depletion results in spine loss associated with synapse elimination**

**(A)** Experimental timeline and constructs used to compare spine dynamics in dTAG-Drebrin-, dTAG-sDrebrin- and GFP-expressing neurons following dTAG-13 treatment and washout. **(B)** Example images of a dendritic branch expressing dTAG-sDrebrin-GFP before (t0) and after (t24) dTAG-13 treatment, together with the corresponding TurboRFP structural marker. Filled and open arrowheads indicate persistent and eliminated spines, respectively. The color bar represents arbitrary fluorescence units. Scale bar = 5 µm. **(C)** Chi-square analysis of persistent and eliminated spine populations in GFP- and dTAG-sDrebrin-expressing neurons following dTAG-13 treatment (t24) and washout (t48). Raw spine counts are shown above each bar. **(D)** Branch-wise spine loss in GFP-, dTAG-Drebrin- and dTAG-sDrebrin-expressing dendritic branches following dTAG-13 treatment (t24) and after washout (t48). GFP: $n = 7$ branches; dTAG-Drebrin: $n = 10$ branches; dTAG-sDrebrin: $n = 7$ branches. **(E)** Spine sizes, normalized to baseline, before dTAG-13 treatment (t0), after treatment (t24), and following washout (t48). The numbers of spines tracked at t0, t24, and t48 were 133, 116, and 114 for GFP, 129, 74, and 60 for dTAG-Drebrin, and 56, 49, and 46 for dTAG-sDrebrin. **(F)** Experimental timeline and constructs used to assess the colocalization of dTAG-Drebrin and PSD95Δ1.2-GFP before and after dTAG-13 treatment. **(G)**

Example images showing dTAG-Drebrin and PSD95Δ1.2-GFP localization before (t0) and after (t24) dTAG-13 treatment. Filled and open arrowheads indicate persistent and eliminated spines, respectively. Color bars represent arbitrary fluorescence units. Scale bar = 5 μm. **(H)** Spine (left) and shaft synapse (right) counts before (t0) and after (t24) dTAG-13 treatment, showing synapses containing PSD95Δ1.2-GFP only, dTAG-Drebrin only, or both proteins. Nearly all spine and shaft synapses contained both markers before and after treatment.

All experiments were performed in rat organotypic hippocampal slice cultures. Graphical data are presented as mean ± SEM except in (C), where total spine counts are shown. Statistical significance was assessed using Chi-square analysis to compare spine fate proportions in (C), linear mixed-effects models (LMM) for branch-wise spine loss (D), and bias-reduced generalized estimating equations (GEE) for spine sizes (E), followed by model-based contrasts with Holm-Bonferroni correction for multiple comparisons. No inferential statistics were performed for the colocalization counts in (H).

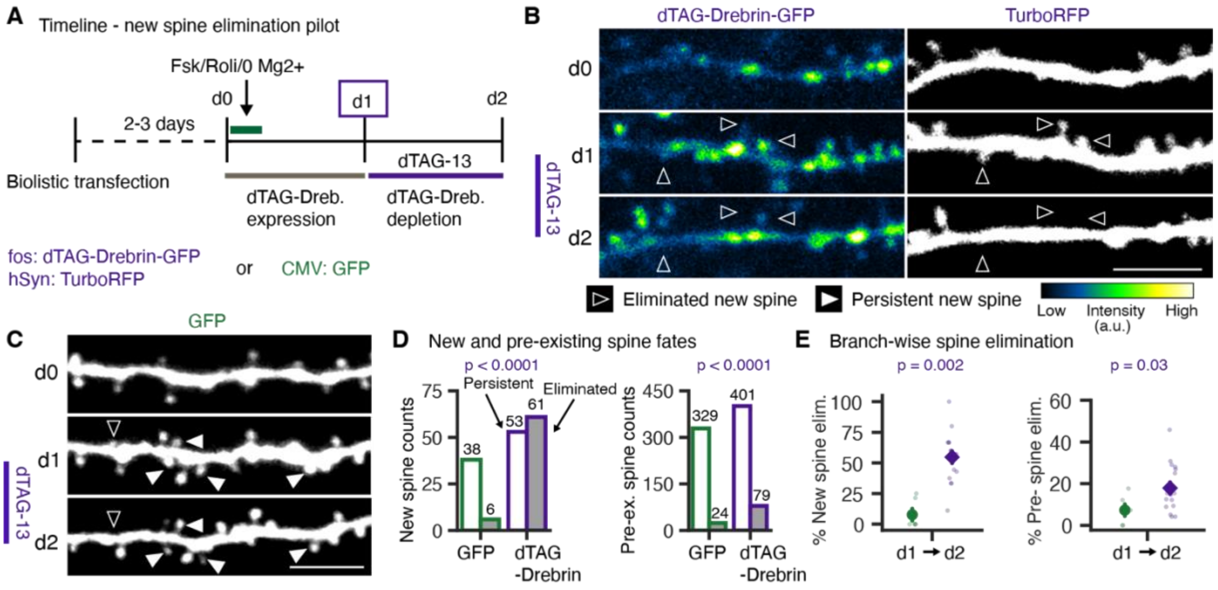


**Fig. S3. Nonselective new spine elimination following activity-dependent, but not doxycycline-dependent dTAG-Drebrin expression**

**(A)** Experimental timeline (top) and constructs (bottom) used for the new spine elimination pilot experiment. Activity-dependent dTAG-Drebrin expression was induced by chemical long-term potentiation (cLTP), and dTAG-13 was administered 24 h later to deplete newly expressed dTAG-Drebrin. **(B, C)** Example images of dendritic branches expressing dTAG-Drebrin-GFP together with the structural marker TurboRFP (B) or GFP (C) before cLTP (d0), after cLTP (d1), and after dTAG-13 treatment (d2). Filled and open arrowheads indicate persistent and eliminated new spines, respectively. Scale bar = 5 µm. Color bar represents arbitrary fluorescence units. **(D)** Chi-square analysis of new (left) and pre-existing spine fates (right) in pooled GFP- and dTAG-Drebrin-expressing spine populations. Raw spine counts are shown above individual bars. **(E)** Branch-wise elimination of newly formed (left) and pre-existing (right) spines following dTAG-13 treatment. GFP: n = 9 branches; dTAG-Drebrin: n = 17 branches. Only branches in which more than two new spines formed following cLTP induction were included in the analysis.

All experiments were performed in rat organotypic hippocampal slice cultures. Graphical data are presented as raw spine counts in (D) and mean ± SEM in (E). Statistical significance was assessed using Chi-square analysis to compare spine fate proportions in (D), and LMM followed by model-based contrasts for branch-wise spine elimination (E).

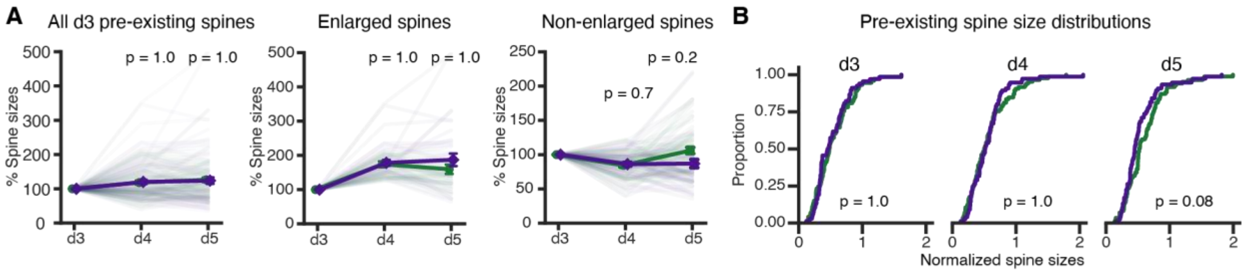


**Fig. S4. Selective new spine elimination is not accompanied by pre-existing spine shrinkage – related to Figure 2**

**(A)** Size dynamics of d3 pre-existing spines in dTAG-Drebrin and "No Dox" control branches, normalized to d3 baseline. Left, all d3 pre-existing spines: 76 dTAG-Drebrin and 85 "No Dox" control spines. Middle, pre-existing spines that enlarged by ≥25% on d4 relative to d3 baseline: 28 dTAG-Drebrin and 32 "No Dox" control spines. Right, remaining pre-existing spines that did not enlarge by ≥25% on d4 relative to d3 baseline: 48 dTAG-Drebrin and 53 "No Dox" control spines. **(B)** Cumulative distributions of normalized spine sizes for d3 pre-existing spines at d3, d4, and d5.

All experiments were performed in Thy1-GFP organotypic mouse hippocampal slice cultures. Graphical data are presented as mean ± SEM. Statistical significance in (A) was assessed using bias-reduced GEE followed by model-based contrasts with Holm-Bonferroni correction for multiple comparisons. Distributions in (B) were compared using two-sample Kolmogorov–Smirnov tests with Holm correction across time points.

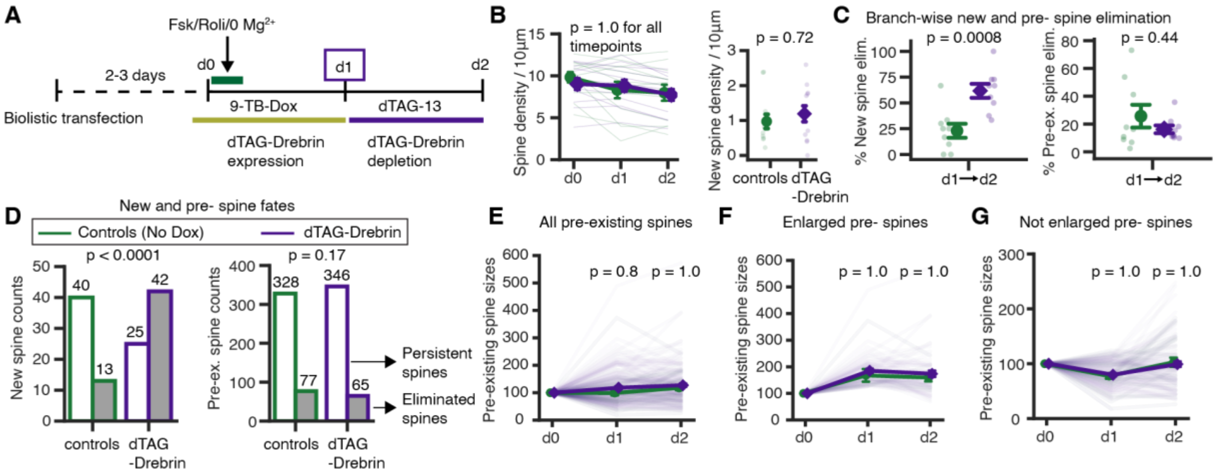


**Fig. S5. Replication of selective new spine elimination following Forskolin/Rolipram-induced structural plasticity**

**(A)** Experimental timeline for a confirmatory selective new spine elimination assay. Structural plasticity was induced using Forskolin/Rolipram in $Mg^{2+}$ -free ACSF. **(B)** Branch-wise total (left) and new spine density (right), normalized to dendritic path length (spines/10 µm). Controls: n = 13 branches. dTAG-Drebrin group: n = 12 branches. **(C)** Branch-wise elimination of newly formed (left) and pre-existing (right) spines in dendritic branches that generated more than two new spines. Controls: n = 9 branches. dTAG-Drebrin group: n = 9 branches. **(D)** Chi-square analysis of new spine fates (left) and pre-existing spine fates (right) in pooled GFP- and dTAG-Drebrin-expressing spine populations. Raw spine counts are shown above the bars. **(E-G)** Size dynamics of d0 pre-existing spines normalized to d0 baseline. E, all pre-existing spines: 125 dTAG-Drebrin and 42 control spines. F, pre-existing spines that enlarged by ≥25% relative to baseline: 51 dTAG-Drebrin and 10 control spines. G, remaining pre-existing spines that did not enlarge by ≥25%: 74 dTAG-Drebrin and 32 control spines.

All experiments were performed in C57BL/6 organotypic mouse hippocampal slice cultures. Graphical data are presented as mean ± SEM except in (D), where total spine counts are shown. Statistical significance was assessed using LMMs for total and new spine density (B) and newly formed-spine elimination (C, left), and a bias-reduced GEE for pre-existing-spine elimination (C, right) and spine sizes (E-G), followed by model-based contrasts with Holm correction.

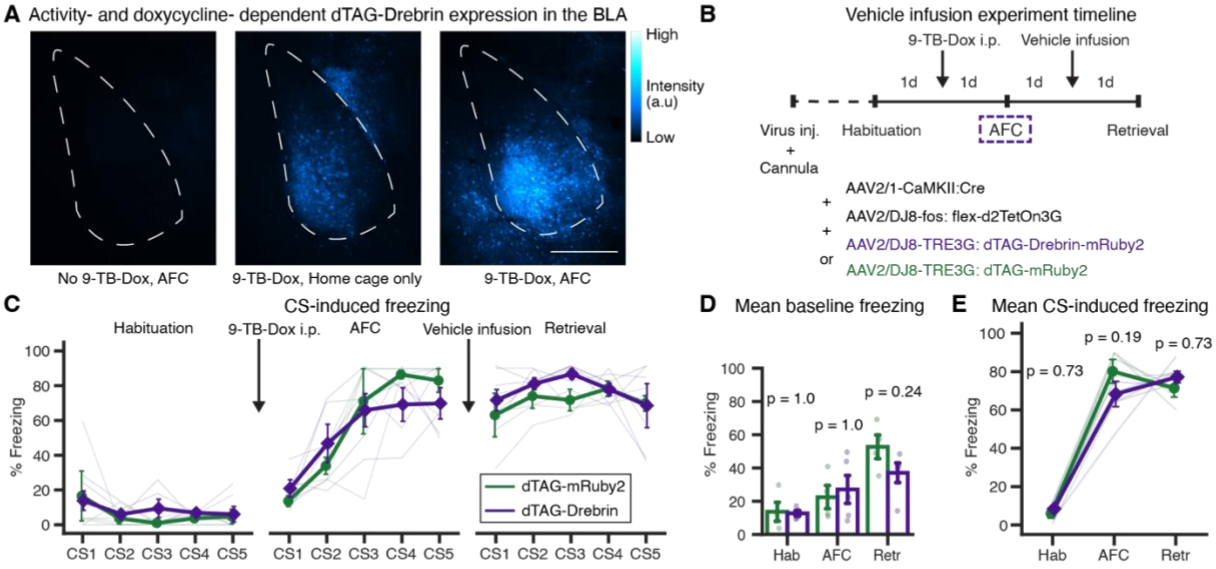


**Fig. S6. Activity- and doxycycline-dependent dTAG-Drebrin expression does not alter auditory fear memory in the absence of dTAG-13 treatment**

**(A)** Verification of activity- and doxycycline (Dox)-dependent dTAG-Drebrin expression in the basolateral amygdala. Example images from mice that did not receive 9-TB-Doxycycline (9-TB-Dox) but underwent auditory fear conditioning (AFC) (left), received 9-TB-Dox without AFC (middle), or received 9-TB-Dox followed by AFC (right). Scale bar = 400 µm. Color bar represents arbitrary fluorescence units. The BLA outline is overlaid on the images. **(B)** Experimental timeline and viral constructs used for the vehicle infusion control experiment. **(C)** Percent freezing to individual conditioned stimulus (CS) presentations during Habituation (Hab), AFC, and Retrieval (Retr) in 4 control and 5 dTAG-Drebrin mice. **(D)** Mean baseline freezing during each behavioral session in the same mice shown in (C). **(E)** Mean CS-induced freezing during each behavioral session in the same mice shown in (C).

Graphical data are presented as mean ± SEM. No inferential statistics were performed for (C), which is included to visualize freezing across individual CS presentations. Statistical significance was assessed using LMM (D) or bias-reduced GEE (E). The p-values shown were obtained from model-based contrasts with Holm–Bonferroni correction for multiple comparisons.

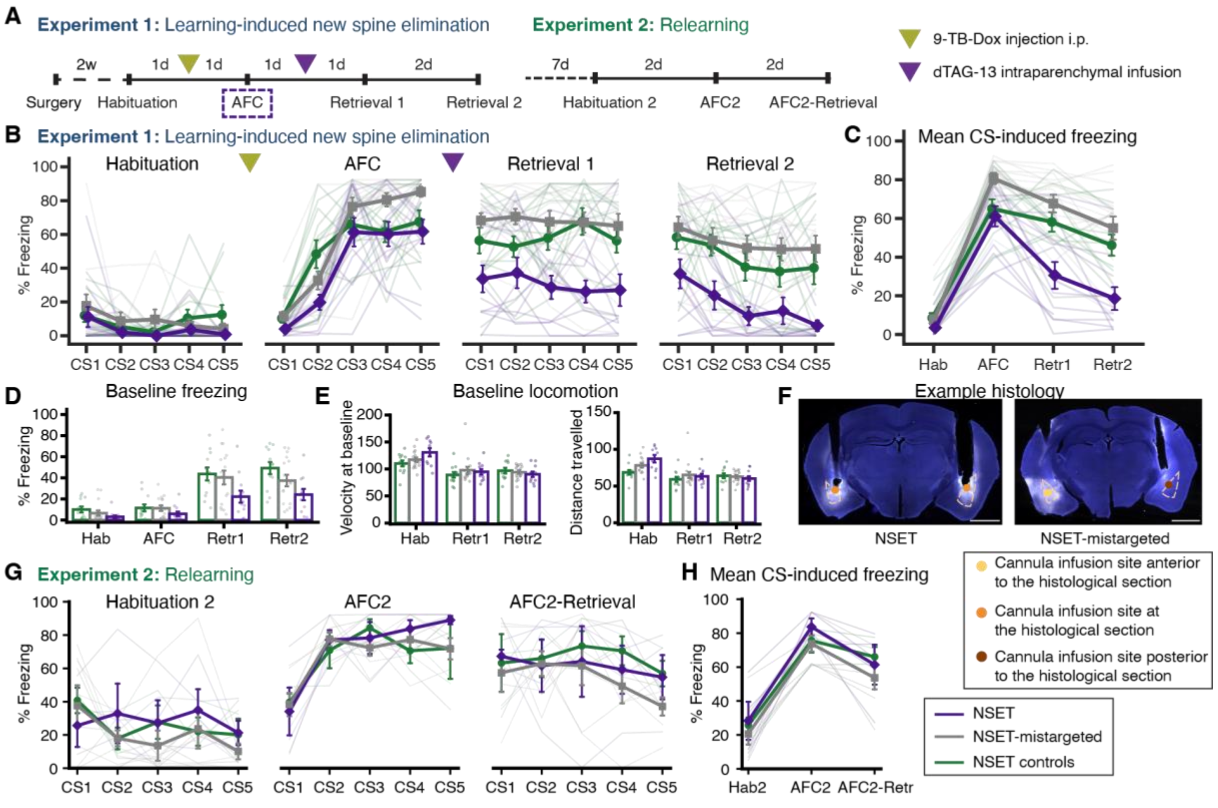


**Fig. S7. Mistargeted NSET mice do not show memory loss**

**(A)** Experimental timelines of the learning-induced spine elimination and relearning paradigms. **(B, C)** Percent freezing during individual CS presentations (B) and mean CS-induced freezing (C) during Habituation, Auditory Fear Conditioning, Retrieval 1, and Retrieval 2 in the learning-induced new spine elimination paradigm. Data are shown for 13 control, 14 NSET-mistargeted and 12 NSET mice. **(D)** Mean baseline freezing during each behavioral session in the same mice shown in (B, C). **(E)** Locomotor activity during the baseline period, excluding freezing epochs, shown as mean velocity (mm/s) and total distance traveled (mm) in the same mice shown in (B-D). **(F)** Example histological sections from an NSET (left) and NSET-mistargeted (right) mouse. Approximate BLA outlines and cannula infusion sites are superimposed on the images. Scale bar = 1.5 mm **(G, H)** Percent freezing during individual CS presentations (G) and mean CS-induced freezing (H) during Habituation 2 (Hab2), Auditory Fear Conditioning 2 (AFC2), and Retrieval following AFC2 (AFC2-Retr) in the relearning paradigm. Data are shown for 3 controls, 7 NSET-mistargeted, and 4 NSET mice.

Graphical data are presented as mean ± SEM. No inferential statistics were performed for any panels in this figure.

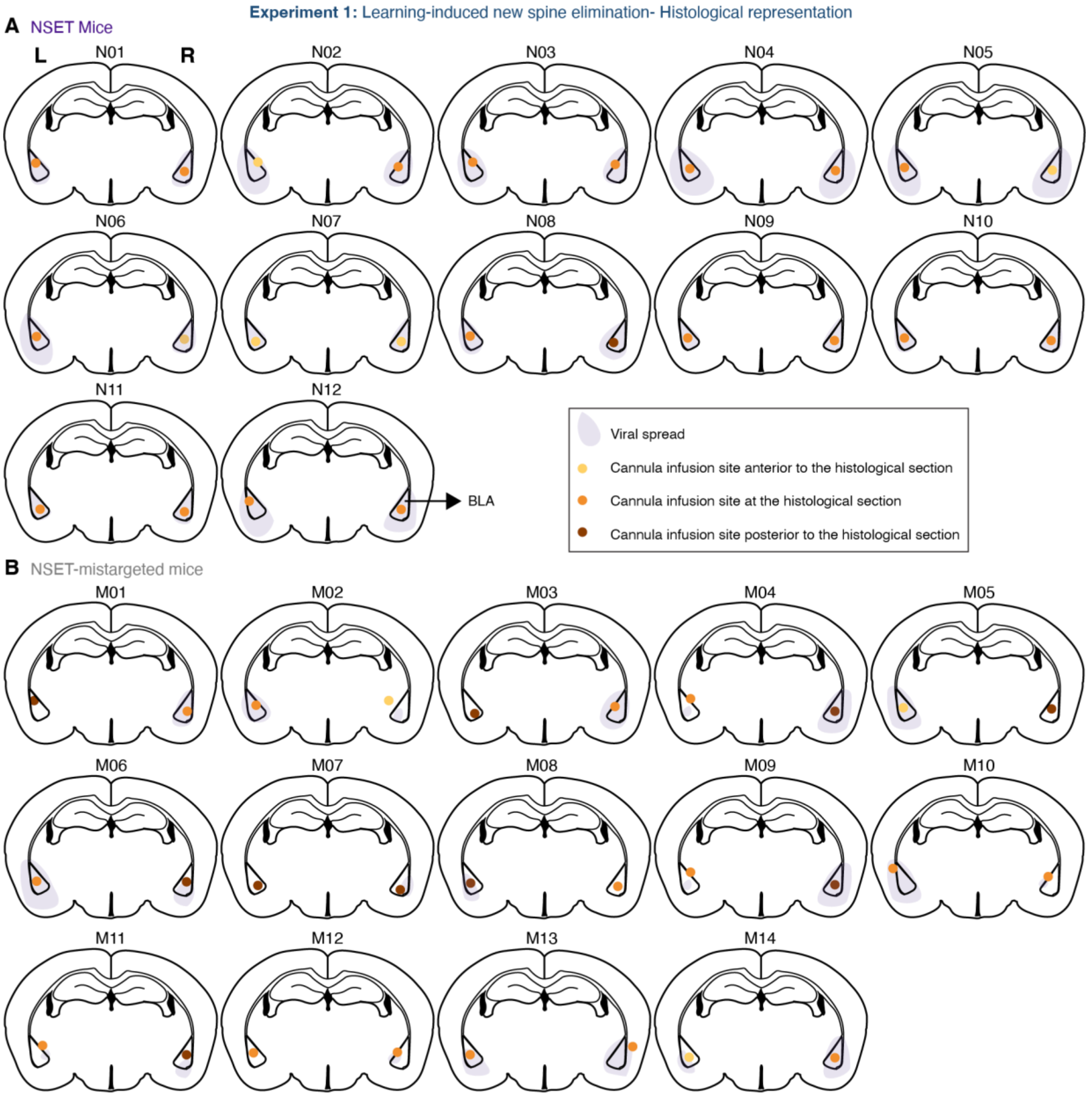


**Fig. S8. Histological representation of NSET and NSET-mistargeted mice from the learning-induced spine elimination experiment**

**(A, B)** Approximate virus spread and cannula infusion site locations relative to the coronal section at bregma −1.84 mm in 12 NSET (A) and 14 NSET-mistargeted mice (B). Shaded regions indicate approximate viral spread; colored points indicate whether the infusion site was anterior to, at, or posterior to the illustrated atlas level.

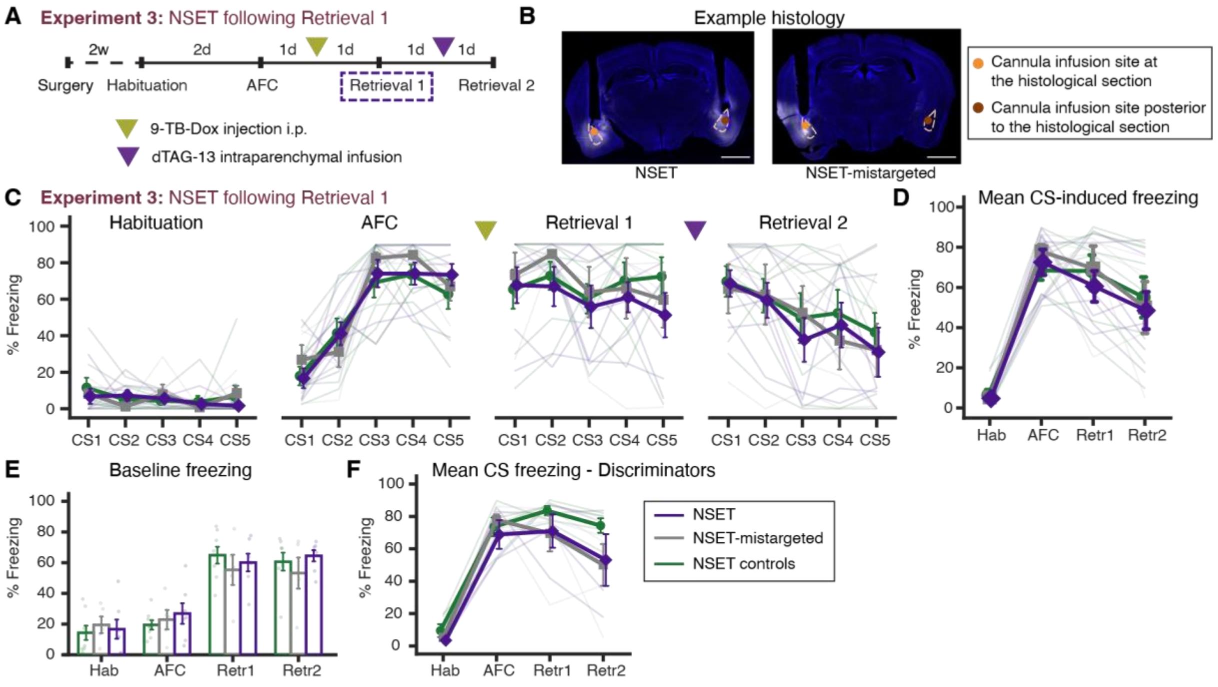


**Fig. S9. NSET application following Retrieval 1 does not affect memory**

**(A)** Experimental timeline for the NSET application following Retrieval 1 experiment. **(B)** Example histological sections from an NSET (left) and NSET-mistargeted (right) mouse. Approximate BLA outlines and cannula infusion sites are superimposed on the images. Scale bar = 1.5 mm. **(C, D)** Percent freezing during individual CS presentations (C) and mean CS-induced freezing (D) during Habituation, Auditory Fear Conditioning, Retrieval 1, and Retrieval 2. Data are shown for 8 control, 5 NSET-mistargeted and 7 NSET mice. **(E)** Mean baseline freezing during each behavioral session in the same mice shown in (C, D). **(F)** Mean CS-induced freezing in "discriminators," defined as mice whose CS-induced freezing during Retrieval 1 exceeded their baseline freezing. Data are shown for 5 control, 5 NSET-mistargeted and 4 NSET mice.

Graphical data are presented as mean ± SEM. No inferential statistics were performed for any panel in this figure.

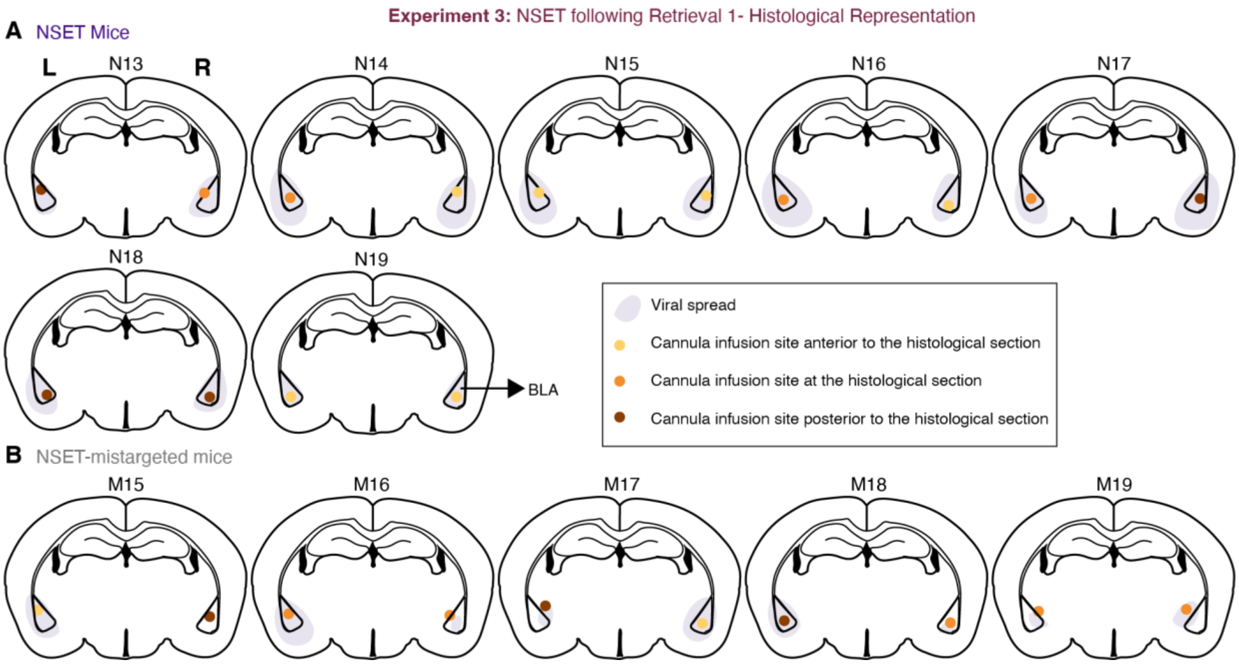


**Fig. S10. Histological representation of NSET and NSET-mistargeted mice from the 'NSET following Retrieval 1' experiment**

**(A, B)** Approximate virus spread and cannula infusion site locations relative to the coronal section at bregma −1.84 mm in 7 NSET (A) and 5 NSET-mistargeted mice (B). Shaded regions indicate approximate viral spread; colored points indicate whether the infusion site was anterior to, at, or posterior to the illustrated atlas level.
